# Crowdsourced RNA design discovers diverse, reversible, efficient, self-contained molecular sensors

**DOI:** 10.1101/2019.12.16.877183

**Authors:** Johan O. L. Andreasson, Michael R. Gotrik, Michelle J. Wu, Hannah K. Wayment-Steele, Wipapat Kladwang, Fernando Portela, Roger Wellington-Oguri, Eterna Participants, Rhiju Das, William J. Greenleaf

## Abstract

Internet-based scientific communities promise a means to apply distributed, diverse human intelligence towards previously intractable scientific problems. However, current implementations have not allowed communities to propose experiments to test all emerging hypotheses at scale or to modify hypotheses in response to experiments. We report high-throughput methods for molecular characterization of nucleic acids that enable the large-scale videogame-based crowdsourcing of functional RNA sensor design, followed by high-throughput functional characterization. Iterative design testing of thousands of crowdsourced RNA sensor designs produced near-thermodynamically optimal and reversible RNA switches that act as self-contained molecular sensors and couple five distinct small molecule inputs to three distinct protein binding and fluorogenic outputs—results that surpass computational and expert-based design. This work represents a new paradigm for widely distributed experimental bioscience.

**One Sentence Summary:** Online community discovers standalone RNA sensors.

## Main Text

A number of recent platforms have demonstrated the engagement, via the internet, of vast untapped human cognitive potential, transforming the scale by which scientific questions can be approached with human intelligence. However, previous work has only allowed “citizen scientists” to process data already collected^1,2^, to interface with purely computational methods to implement solution strategies^3^, or to compete for limited slots on a low throughput experimental synthesis pipeline^4–6^. We reasoned that by enabling a community of RNA design enthusiasts to both generate hypotheses and then acquire large-scale experimental evidence supporting or rejecting these hypotheses, we could “close the loop” on crowd-sourced science, providing a platform for iterative hypothesis generation and experimental testing. Recent advances in high-throughput functional characterization of nucleic acids on sequencing instruments provides potentially immense capacity for carrying out experiments devised by these scientific communities^7–9^. We hypothesized that with these tools, online communities would be capable of producing RNA sensors with unprecedented performance as well as principles that might generalize to other unsolved scientific design tasks.

RNA is an attractive substrate for diagnostics^10^, cellular computing^11^, ‘smart’ imaging tools^12^ and targeted therapeutics^13,14^ due to its inherent biocompatibility, capacity for large-scale structural rearrangements via alternative base pairings, and ability to form aptamers that specifically recognize cellular metabolites, signaling molecules, ions, cofactors, and proteins^15,16^. Such ‘riboswitches’ sense biomolecules and respond by causing RNA cleavage^17^, transcription termination^18^, or other altered genetic outputs. However, these thermodynamically irreversible output signals are not compatible with near real-time measurements from self-contained and reversible switches, and thus preclude important applications, such as fluorescence-based sensors of temporally dynamic processes or reversible chemical control of therapeutics^19^. To date, all ‘standalone’ riboswitches have exhibited activation ratios (AR, the ratio of the measured output signal in the presence versus absence of ligand) of less than 10-fold—far below the limits allowed by thermodynamics^20^—and a thermodynamically reversible RNA switch has not been demonstrated. Thus, a method to design efficient, standalone RNAs capable of conformationally responding to inputs both sensitively and reversibly is needed. This outstanding challenge in RNA design provides an ideal testbed for distributed, internet-based science.

To implement a platform for hypothesis-based, iterative scientific discovery for RNA sensors, we combined internet-scale crowdsourcing of RNA design using the scientific discovery game Eterna^55^ and high-throughput, quantitative RNA characterization using repurposed sequencing chips and instruments^21,22^. Eterna puzzles challenge an online player community to design RNA sequences that couple the formation of a ligand-binding input element (**Fig. 1A**) to promote or prevent the formation of a second, sequence-defined output element. These output elements provide measurable fluorescent readout proportional to a switch’s response. The Eterna platform enables players to design sequences that will fold differentially in the presence of ligand, using standard thermodynamic modeling packages to estimate free energies of RNA folding and to simulate binding of the ligand. While providing rapid computational feedback, these packages provide imperfect approximations of the expected folding of model designs^23^, necessitating both human insight and experimental testing. We solicited tens of thousands of Eterna designs from the online community (**Fig. 1B**), and then characterized these designs using massively-parallel RNA array technology. Briefly, player designs were displayed on Illumina sequencing chips and switch behavior was evaluated under relevant conditions (RNA-MaP, RNA on a massively parallel array; **Fig. 1C–E**). This platform quantifies switch function by measuring binding curves for every design interacting with a fluorescent reporter molecule in both the presence and absence of the input ligand (**Fig. 1F**) and achieves excellent reproducibility (*R*^2^ = 0.94; **Fig. S1**)^23–25^. Results from these experiments were then released to the Eterna community, and further design refinement was solicited (**Fig. 1A**).

**Fig. 1.**
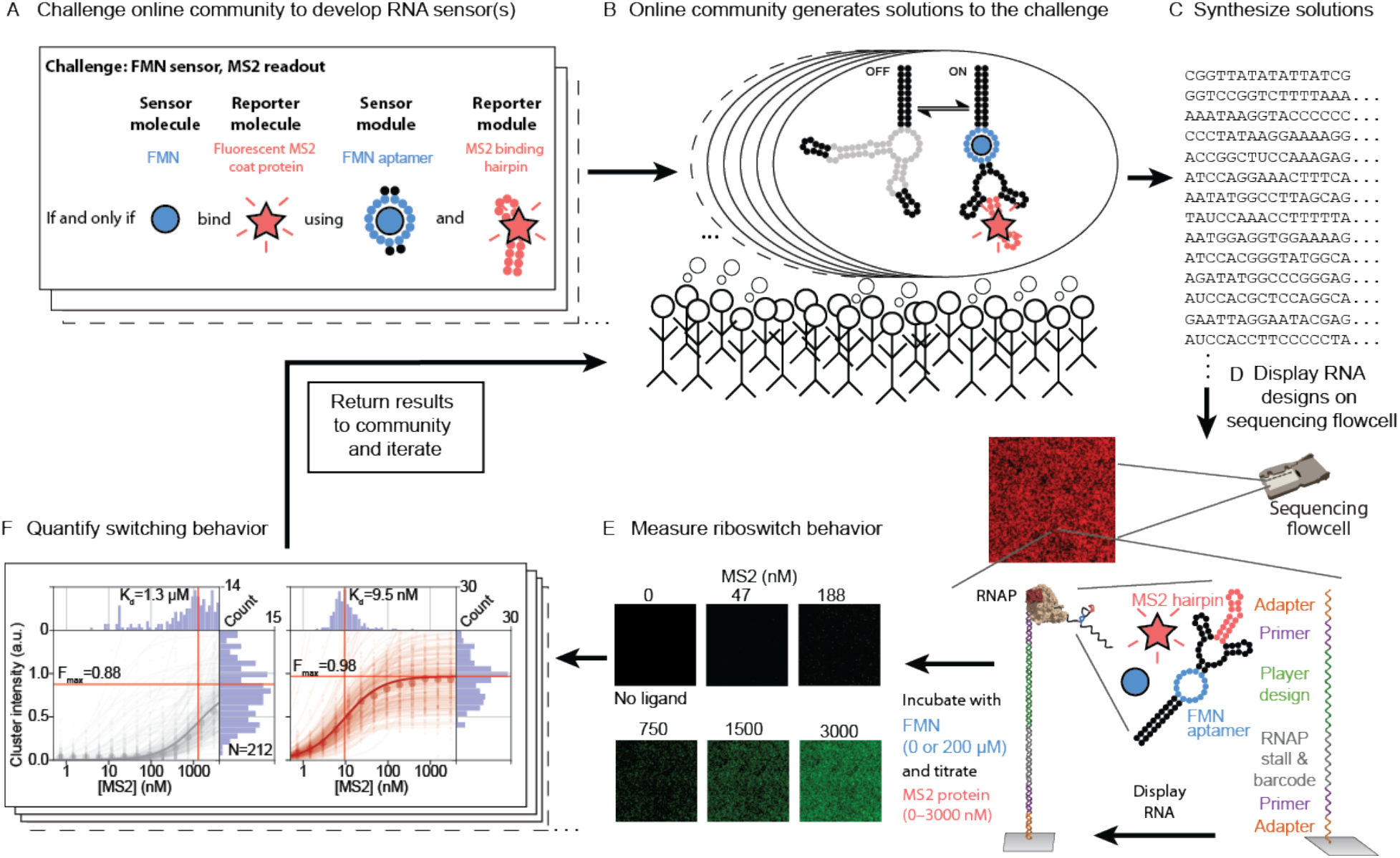
A platform for crowdsourced RNA sensor design and testing at high-throughput. A) Riboswitch sensor design challenges are released to the Eterna community. In this example, an FMN aptamer (small blue circles) should be used to sense the FMN molecule (large blue circle), and in the presence of FMN an MS2 hairpin (red circles) must fold for detection via binding of fluorescent MS2 coat protein (red star). B) Eterna community designs sequences of riboswitches predicted to fold into appropriate OFF and ON states. C) Puzzle solutions are synthesized by DNA array synthesis and converted to libraries ready for RNA-MaP (RNA on a massively parallel array) via PCR. D) Following high-throughput sequencing of the library on an Illumina sequencing chip, a biotinylated primer is annealed and extended to ssDNA. Streptavidin is then bound and RNA polymerase displays one riboswitch variant solution per sequencing cluster. E) Binding of fluorescently labeled MS2 coat protein is then quantified across all clusters at increasing concentrations (example images shown). Cluster sequence information is used to link fluorescent signals to underlying riboswitch sequence. F) Binding data is quantified from multiple clusters for a single riboswitch variant (median fits shown as bold line). These data are then released to the Eterna community and subsequent rounds of designs are solicited.

As a first test, we challenged players to couple input and output RNA aptamers that were individually well-characterized but which had not been previously combined into a riboswitch: an 11-nt aptamer for the cellular metabolite flavin mononucleotide (FMN; input)^26^ and a 19-nt hairpin that binds the MS2 coat protein (output)^21^. The maximum activation ratio (*AR*_max_) expected for this challenge, as determined by the dissociation constant of the FMN aptamer and the experimental trigger concentration of input ligand, was 132±33 ^20^ (Supplemental Text). Over seven consecutive rounds of design, synthesis, characterization, and community-wide discussion, 149 members of the Eterna community submitted over 30,000 designs (**Fig. S1**). Of these, 30 designs from 5 distinct individuals fell within error of the thermodynamic maximum (**Fig. 2A; Fig. S2; Table S2**).

**Fig. 2.**
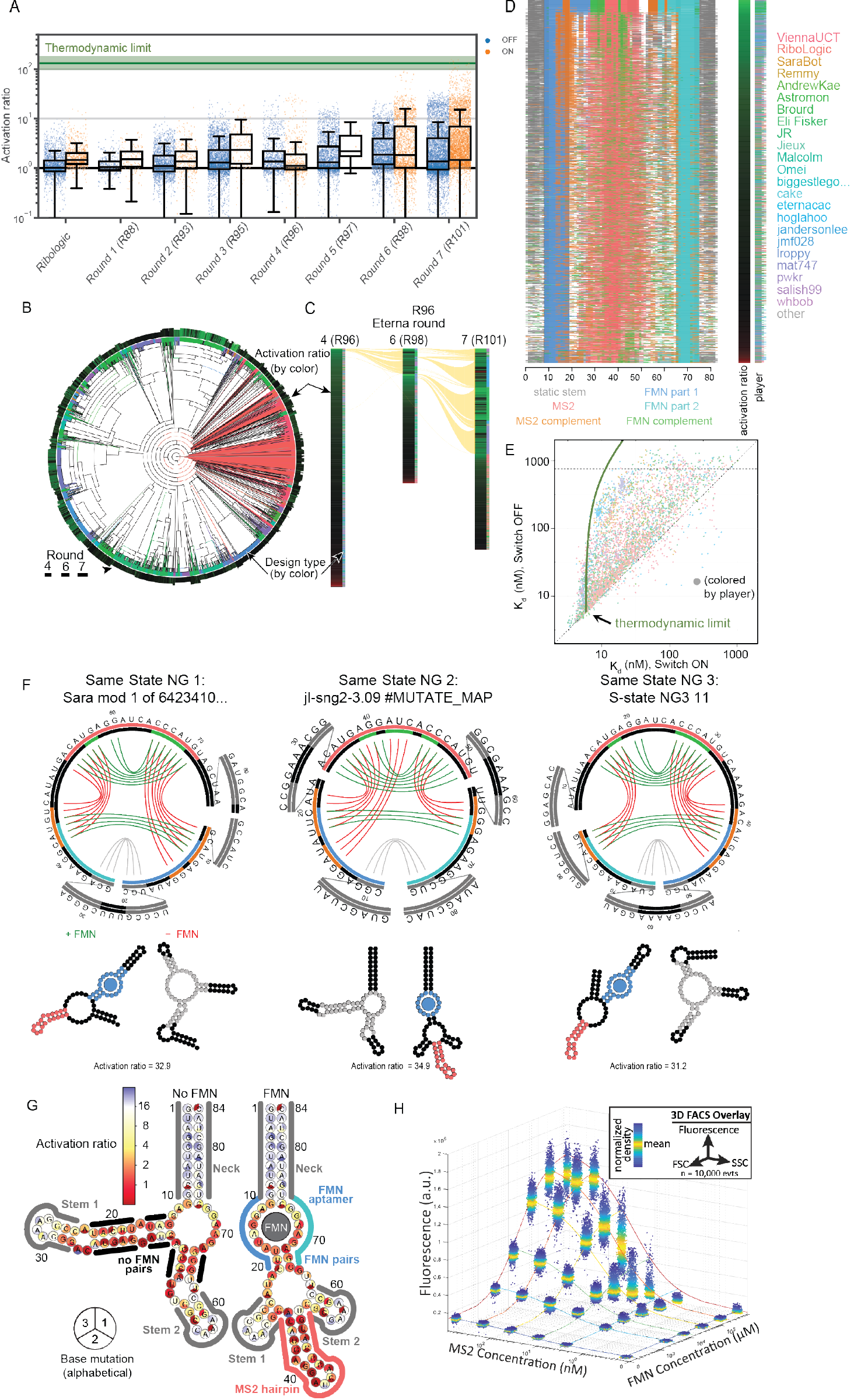
Performance of player-designed sensors. A) Activation ratio for ON (blue points) and OFF (orange points) FMN-MS2 switches over 7 rounds show substantial improvement as players learn from prior rounds of design and outperform automated Ribologic design. The theoretical thermodynamic limit for activation ratio, estimated using the measured intrinsic *K*_d_ of the FMN aptamer, is depicted as a green line. Error bars represent propagated standard error from *K*_d_ estimation. B) Phylogenetic tree of puzzle solutions for an FMN-MS2 ON switch (“Same State NG 2”). Terminal branches are colored by design type: player designs (green), automated solutions from Ribologic (red), and modifications of previous designs (blue). The outer ring is colored by activation ratio (brighter green is higher, red is less than 1) and the width represents the round of submission (1–3). C) “Same State NG 2” solutions by round, color-coded as in B). Yellow lines connect “mods,” defined as sequences separated by Levenshtein edit distance <5. D) Position of functional elements across all “Same State NG 2” solutions, color-coded as indicated, sorted by activation ratio. The MS2 and FMN complements are defined as parts of the sequence predicted to pair with any part of the MS2 hairpin or FMN aptamer, respectively. E) *K*_d_ values for both states of “Same State NG 2” puzzle designs. The predicted thermodynamic limit for the *K*_d_ in the OFF-state, based on the *K*_d_ in the ON-state, is shown in green. F) Predicted secondary structures for the best solutions from three ON-switch puzzles with predicted invariant base pairs (grey segments) and base pairs in the absence (red) or presence (green) of the FMN ligand. Outer circles display the MS2 hairpin and complementary segments in color; inner circle displays the FMN aptamer and complementary segments. G) Activation ratio for switching by mutants of the top-scoring design, mapped by color onto predicted secondary structures. Blue implies less disruption of switching upon mutation. Perturbations of single mutations for each base are indicated by color within circle sectors (A, C, G, or U proceeding clockwise from top and omitting the original base). Sequence segments representing functional (FMN in blue/turquoise, MS2 in red), invariant (grey) or other secondary structures elements are outlined by color. H) Fluorescence intensity of particle display beads under varying concentrations of FMN and MS2. Each cluster of points is a 3D overlay of a scatter plot showing the measured forward scatter (FSC), side scatter (SSC), and fluorescence intensity of an RNA-coated particle centered at a given FMN and MS2 concentration. Colored lines represent the global fit of the mean fluorescent values at each concentration to an equilibrium binding model.

In community-wide discussions, players attributed their success to two primary factors: (1) the relaxation of puzzle design constraints (enabled through updates of the Eterna source code) that allowed exploration of a much larger design space, and (2) the public dissemination of annotated experimental data that let players iterate and improve upon any past design. Designs exhibited striking improvements over iterative rounds; in the first round, the maximum AR value achieved was 5.7, while by the last rounds several designs exhibited AR values in excess of 80. The most dramatic improvement occurred after the fourth round when constraints on the ordering of the FMN and MS2 hairpin were eliminated (**Fig. S1, Table S1**). This game update allowed the community to autonomously explore a wide variety of such element orderings (i.e., tandem or split-aptamer arrangements) in remaining rounds (represented as clades in **Fig. 2B**).

Changes to design constraints and iterative refinement of promising solutions (**Fig. 2C**) yielded a collection of structural mechanisms (i.e., orderings of aptamer elements and design of sites reverse complementary to those elements, **Fig. 2D**) that reliably yielded high *AR* values, often approaching the thermodynamic maximum AR_max_ of 132 (green bar, **Fig. 2A**). As expected from theoretical predictions^20^, these top designs exchange lower output levels in the activated (reporter-bound) state in exchange for a higher overall activation ratio (**Fig. 2E**). The relaxation of design constraints also allowed the design of double FMN aptamer switches, a “cheat” strategy that allows designs to exhibit *AR* values above the thermodynamic limit for a single FMN-bound switch (**Fig. S3**).

Computational predictions of RNA secondary structures for top-performing switches revealed the particular utility of interleaving base pairs between the FMN and MS2 aptamers in the inactive state (misfolded reporter) and of ‘miniaturizing’ designs by promoting stable base pairings between the 5′ and 3′ regions or non-switching stems and hairpins that sequester non-switching nucleotides (**Figs. 2B,F**). To validate these structural mechanisms, we carried out SHAPE chemical mapping^27^ and single mutant as well as compensatory rescue^28^ analysis for a top-scoring design “JL-sng2-3.09”; these data confirmed the predicted secondary structures in both ligand-free and ligand-bound states (**Fig. 2G** and **Fig. S4**). To further test the general usability of these sensors in different experimental contexts, we shifted our readout from RNA-MaP to cytometric analysis of riboswitches displayed on micro-magnetic particles^29^. Using this orthogonal assay, “JL-sng2-3.09” exhibits robust switching behavior across a wide dynamic range of both FMN and MS2 concentrations, suggesting these designs are broadly transferrable to other assays (**Fig. 2H**).

The impressive performance of player-designed switches encouraged us to compare a concurrently developed, physics-based computational algorithm for riboswitch design—Ribologic^19^—against the performance of the Eterna community. In the FMN sensor challenges, Ribologic designs gave lower activation ratios than the final player designs after iterative refinement (**Fig. 2A**). To compare Ribologic designs to player-curated designs prior to iterative refinement, we performed an additional round of MS2-activated small molecule switches using RNA-MaP technology. Here, we introduced new small molecule aptamers that bind theophylline and L-tryptophan and challenged players and Ribologic to design diverse ON and OFF switches. In each of these 8 puzzles, the best crowd-sourced designs outperformed Ribologic-generated designs by consistently achieving *AR* values above 10 and, in several cases, approaching the thermodynamically allowed maximum (**Fig. 3A, Fig. S2**).

**Fig. 3.**
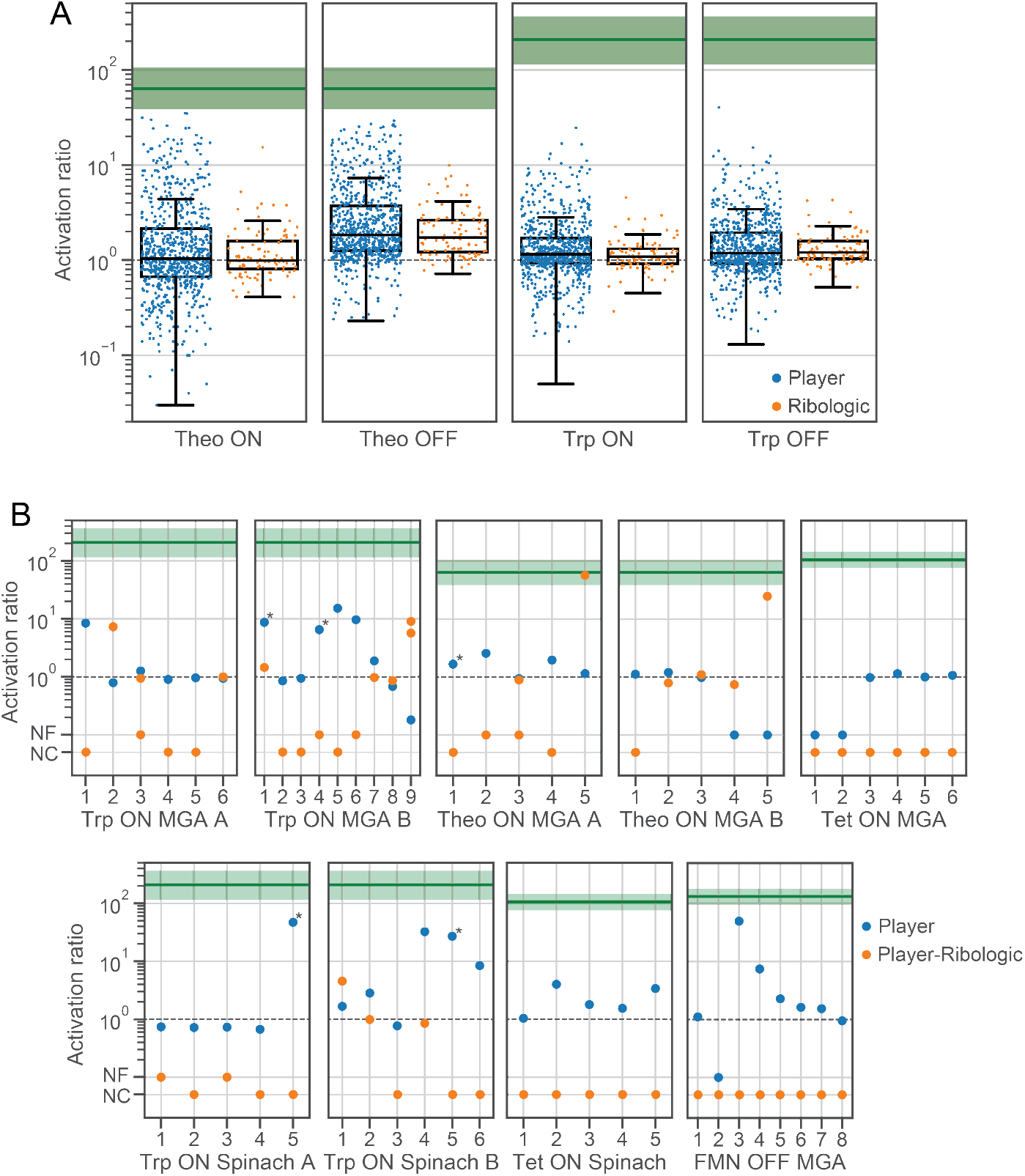
Single-round design challenges on alternative inputs and output. (A) Player designs of both ON and OFF switches coupling the theophylline (Theo) and tryptophan (Trp) aptamers to MS2 coat protein binding exhibited significantly higher activation ratios than Ribologic-designed solutions. Theophylline solutions approached the thermodynamic limit for activation ratios (depicted as green line, estimated from measured intrinsic ligand aptamer *K*_d_ values). (B) “Light-up” round designs for tryptophan (Trp), theophylline (Theo), tetracycline (Tet), and FMN input ligands coupled to folding of malachite green (MGA) or Spinach light-up aptamers. Each vertical line represents a different switch architecture, designed by a player, that was used as a starting point for the corresponding Ribologic design in the same column. Some player designs were used to seed more than one Ribologic solution. NF, the tested design was nonfunctional in both states (no detectable binding to fluorophore); NC, implies that Ribologic was unable to converge to a solution for the given design; ‘*’ marks designs confirmed across multiple independent measurements, with the mean activation ratio displayed.

Next, we asked if players could apply the lessons learned from past rounds to produce winning solutions when given dramatically fewer overall experimental attempts. To prevent modifications of past successful designs, we introduced two new reporting aptamers—the malachite green aptamer (MGA)^30,31^ and the DFHBI-binding aptamer RNA Spinach^32^ – which bind to and activate the fluorescence of otherwise non-fluorescent small molecule dyes – and designed puzzles that coupled small molecule aptamers against tryptophan, theophylline, and FMN, as well as, for the first time, tetracycline^33^. For these “light-up” challenges, we reduced the number of tested designs to fewer than 10 per puzzle, solicited solutions through bi-weekly community-wide voting, and used solution-phase fluorescence as a new readout. Despite having dramatically fewer attempts (only 72 designs were tested over all light-up challenges), players continued to deliver highly responsive switches (**Fig. 3B**) with several *AR* values exceeding 15. Confirming the difficulty of these challenges, attempts to use Ribologic to generate computational solutions failed, due to the large search space involved in positioning sensor and reporter elements.

We next asked if player guidance could improve the automated design of riboswitches. We then used player solutions to seed the designs for player-Ribologic hybrid designs. Fixing the location of the included aptamer motifs allowed Ribologic to converge on new designs based on player-submitted solutions. In two examples, coupling player-guided aptamer placement with the Ribologic design infrastructure yielded switches surpassing the best player designs, suggesting that experience-guided aptamer and reporter placement may enable semi-automated design of highly functional molecular sensors, and suggesting new routes to approach next-generation riboswitch design algorithms. Further analysis of the features of the most successful designs is presented in **Supplementary Text**.

An important and previously undemonstrated goal for RNA design is the reusability of designed RNA switches to enable energy-neutral applications. In our final experiments, we therefore tested the reversibility of switches by exposing 482 FMN-MS2 Eterna designs to 52 buffer exchanges (cycling between 0 and 200 μM FMN) over 29 h. In all switches, we observed consistent toggling of the fluorescence output (**Figs. 4A,B**) with many designs exhibiting only minor functional degradation over the length of the assay—suggesting near full reversibility. To test switching in the absence of the MS2 reporter molecule, we immobilized the “JL-sng2-3.09” design on DNA-coated beads and carried out 80 FMN buffer exchanges, subjecting aliquots at each point to dimethyl sulfate (DMS) chemical mapping to assess RNA conformational changes. We observed toggling of the global RNA structure through the entire time course (**Figs. 4C–E**). Finally, we measured the kinetics of activation for two tryptophan-responsive light-up sensors coupled to the MGA and Spinach aptamers. Both yielded design-specific responses to their respective ligands on the order of seconds to minutes, confirming their suitability for kinetic measurements on the timescale of tens of minutes or better (**Fig. S5**).

**Fig. 4.**
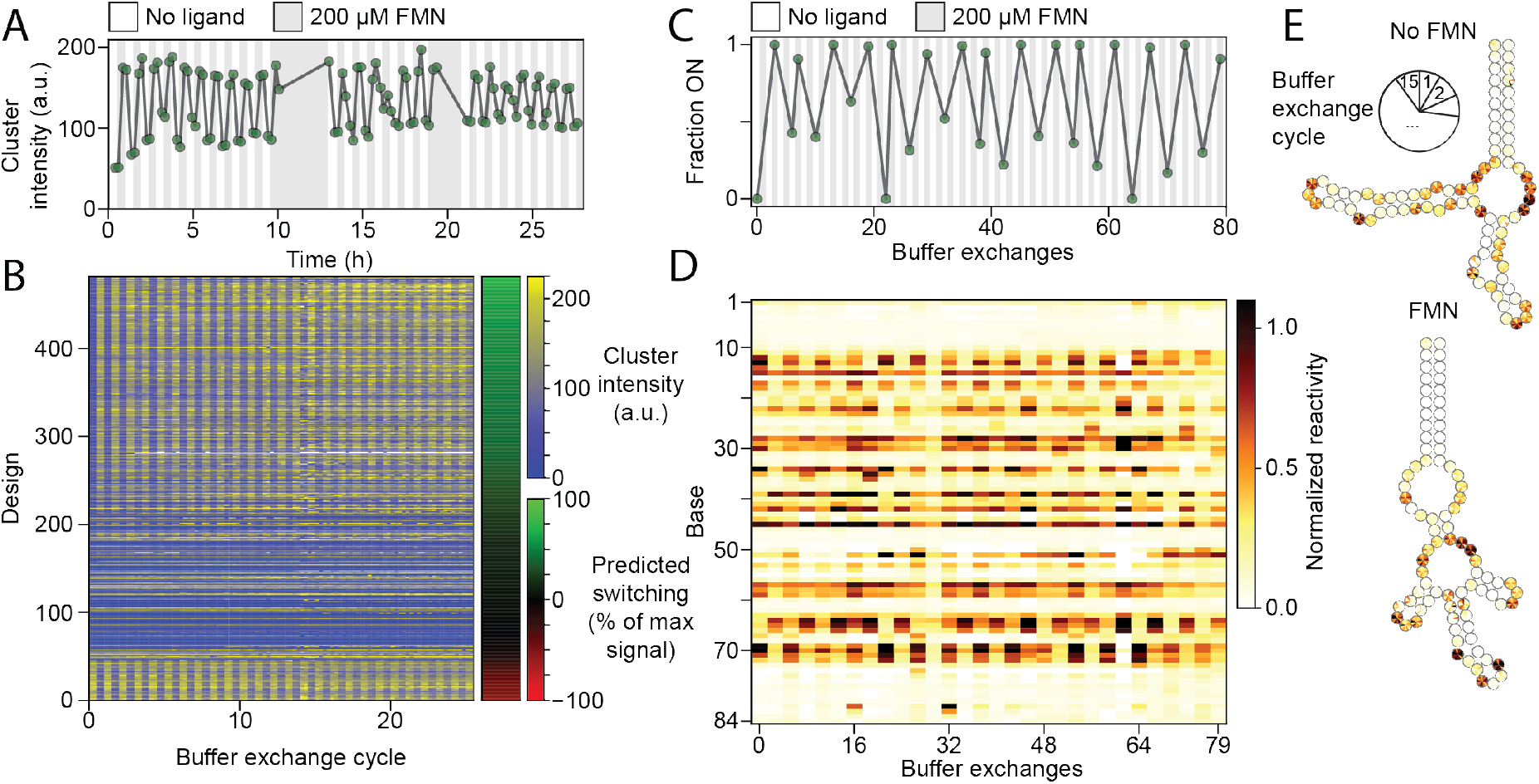
Reversible switching of riboswitches. A) Fluorescence signal from top-scoring single-aptamer design as FMN concentration is repeatedly toggled from 0 (white) to 200 μM (grey). Signals from up to 112 individual clusters were averaged. B) FMN switches from the same experiment in panel A, ordered by predicted switching magnitude and direction (right bar). Bottom constructs are predicted to switch in the opposite direction (red color in the right bar). C) Switch behavior across 80 buffer exchanges as probed by chemical mapping (dimethyl sulfate probing carried out on aliquots after every third exchange). D) Reactivity per base for aptamer profiled in C). E) Reactivity per base mapped onto predicted secondary structures. Each base is divided into 15 segments corresponding to each time point.

This study demonstrates that an iterative, massive open laboratory approach—internet-scale crowdsourcing of experimental hypotheses in combination with a high-throughput, quantitative experimental pipeline for hypothesis testing—enables design of near optimally efficient, reversible, self-contained riboswitches that couple diverse input molecules to diverse output fluorescence modalities, opening the door to a variety of applications. Importantly, this community-based approach to science demonstrates a general insight: non-classically trained citizen scientists can produce remarkable scientific insights when freely allowed to refine their hypotheses as dictated by the scientific method. We believe other citizen science projects would benefit by enabling their communities to define and test all incoming hypotheses^6^. More specific to the field of synthetic RNA biology, we found it critical to explore diverse motif orderings and structure-toggling mechanisms to achieve optimal performance. The merits of such exploration have not been broadly appreciated by the RNA design field and are likely applicable to other unsolved problems in synthetic biology. Furthermore, we are optimistic that the ongoing revolution in high-throughput biology will engender the creation of other platforms “democratizing” experimental biological science through the provisioning of low-cost experimental validation of hypotheses generated by diverse non-experts.

## Acknowledgments

We thank Jeehyung Lee and Adrien Treuille for early work on the Eterna riboswitch design interface; Eli Fisker and Jeff Anderson-Lee for leading discussions of riboswitch experiments amongst Eterna participants; and Curtis Layton, Caleb Geniesse, Benjamin Keep, F. Portela, Siqi Tian, and John Nicol for design and maintenance of the software and experimental platforms.

## Author contributions

Conceptualization, J.A., M.G., M.W., R.W., W.G., R.D., and Eterna participants; Methodology, J.A., M.G., M.W., F.P., W.K., R.D., W.G.; Software, J.A., M.G., M.W., H.W., F.P., R.D.; Formal Analysis, J.A., M.W., W.K., M.G., R.W., R.D.; Investigation, J.A., W.P., M.G., F.P., R.W., Eterna participants; Writing – Original Draft, J.A.; Writing – Review & Editing, J.A., M.G., M.W., R.D, W.G., Eterna participants; Visualization, J.A., M.G., M.W., H.W.; Supervision, Project Administration, and Funding Acquisition, W.G., R.D.;

## Data and materials availability

The Eterna results are available on the Eterna website. Raw data, analysis code, and materials are available upon request.

## Supplementary Materials

Materials and Methods

Supplemental Figures S1-S5

Supplemental Tables S1-S3

Supplemental File

