## Supplemental material for "Crowdsourced RNA design discovers diverse, reversible, efficient, self-contained molecular sensors"

**This PDF file includes:**

Materials and Methods
Supplementary Text
Figs. S1 to S5
Tables S1 to S3

### Materials and Methods

#### Eterna Online interface

RNA molecules were designed in Eterna (https://eternagame.org), an online platform where participants solve design challenges, or puzzles, directly in a web browser[1]. The game-like interface was updated with improved design and features that help participants, such as indicators for design constraints and a 'stamper' tool that allows for quick interactive entry of predefined sequences, such as the MS2 RNA hairpin. The range of available RNA folding engines was expanded from ViennaRNA 1.8.4[2] to also include ViennaRNA package 2.1.9[3] and NuPACK 3.0.4[4], and participants could toggle between these engines during design. Designs were not required to fold or switch correctly in any of these computational folding engines to be submitted by participants.

#### High-throughput analysis of participant-designed switches by RNA-MaP

Sequencing libraries

For RNA-MaP high-throughput functional assays, Eterna participants' solution sequences were collected in a series of rounds. Within the global round numberings across all Eterna challenges, sequences for this study were collected in R88 to R107; see Fig. S1 and Tables S1-S2. These sequences were prepended and appended with common flanking sequences (see below) and synthesized (CustomArray, Bothell, WA), using the procedure recently described[5]. Briefly, sequencing libraries were prepared from the oligo pool either by PCR or emulsion PCR (ePCR)[6,7] in multiple steps, as follows. First, the oligo pool was amplified with primers Read2 and RNAPstall or RNAPstall2. In the second step, sequences for Illumina sequencing adapters (C_adapter and D_adapter), sequencing primers (Read1 and Read2), RNAP promoter and stall sequences, and, for later Eterna rounds, a 16-nt barcode sequence, were added by PCR. Concentrations were 1.5 nM for the previously amplified oligo pool, 3.8 nM for the flanking DNA oligos (C_Read1_RNAPstall and D_Read2) and 137 nM for outer primers (C_adapter and D_adapter). The optimal number of cycles was determined by gel electrophoresis. For libraries prepared by ePCR, the oligo pool was amplified in one step, as previously described[7].

For libraries with barcodes, a bottlenecking step was performed. The library was diluted to ~700,000 molecules and amplified for 16 cycles with primers C_adapter and D_adapter . Libraries were quantified by qPCR or Qubit (Thermo Fisher Scientific) and sequenced on an Illumina MiSeq sequencer with 150-cycle kits.

#### Oligonucleotide sequences for RNA display

| Primer name | Sequence |
|---|---|
| Read2 | CGGCATTCCTGCTGAACCGCTCTTCCGATCT |
| RNAPstall | GTAAGGAGGTTGTATGGAAGACGTTCCTGGATCC |
| RNAPstall2 | GTAAGGAGGTTGTATGGAAGACGTTCCTGGAT |
| C_adapter | AATGATACGGCGACCACCGAGATCTACAC |
| D_adapter | ATCTCGTATGCCGTCTTCTGCTTG |
| C_Read1_RNAPstall | AATGATACGGCGACCACCGAGATCTACACTCTTTCCCTACACGACGCTCTTCCGATCTTTTATGCTATAATTATTTCATGTAGTAAGGAGGTTGTATGGAAGACGTTCCTGGATCC |
| C_Read1_RNAPstall2 | AATGATACGGCGACCACCGAGATCTACACTCTTTCCCTACACGACGCTCTTCCGATCTTTTATGCTATAATTATTTCATGTAGTAAGGAGGTTGTATGGAAGACGTTCCTGGAT |
| C_Read1_BC_RNAPstall | AATGATACGGCGACCACCGAGATCTACACTCTTTCCCTACACGACGCTCTTCCGATCTNNNNNNNNNNNNNNNNTTTATGCTATAATTATTTCATGTAGTAAGGAGGTTGTATGGAAGACGTTCCTGGAT |

| | |
|---|---|
| D_Read2 | CAAGCAGAAGACGGCATACGAGATCGGTCTCGGCATTCCTGCTGAACCGCTCTTCCGATCT |

Preparation and measurement of riboswitches on an Illumina sequencing chip

Sequencing chips were recovered and mounted on a custom automated fluorescent microscope, as previously described[8]. The binding of a SNAP-549-labeled MS2 coat protein (MS2-dlFG mutant) to each RNA design, in the absence or presence of its target small molecule was measured directly on the flow cell, similar to our previous studies[9]. Briefly, the sequencing chip was recovered and mounted onto a custom sequencer-based microscope where dsDNA templates were generated from the clonal sequencing clusters. Clusters of clonal ssDNA strands were recovered by stripping residual fluorophores and complementary DNA after sequencing. A biotinylated oligonucleotide was hybridized to the ssDNA, and dsDNA templates were generated by Klenow extension. Common sequences were further blocked and streptavidin was added to generate a roadblock at the end of the template. *E. coli* RNAP was initiated and stalled by nucleotide starvation, excess polymerase was removed, and the full design was then generated at saturating nucleotide concentrations for 10 min. The common RNA sequences representing the RNAP stall sequence and the Read2 primer were blocked by hybridizing Cy5-labeled and unlabeled DNA oligonucleotides, respectively, at concentrations of 500 nM during transcription and then in buffer for an additional 10 min. MS2 protein was introduced to the flow cell, starting at 0.7 or 1.5 nM, and increased by factors of two up to 3 µM. Incubation times varied from 0.8–1.5 h at the lowest concentrations to 10–20 min at the highest concentrations. The buffer conditions for RNA-MaP titrations were 100 mM Tris-HCl, pH 7.5, 80 mM KCl, 4 mM $MgCl_2$, 0.1 mg/mL BSA, 1 mM DTT, 0.01 mg/mL yeast tRNA, 0.01% Tween-20. Subsequently, the DNA was stripped, RNA regenerated for each cluster, and the MS2 titration repeated in the presence of constant amounts of each small molecule ligand. Fluorescence images of the generated RNA and the bound MS2 were collected using red (660 nm, 200 mW, 400 ms exposure time) and green lasers (532, 200 mW, 400 ms or 1.0 s), respectively. Each design was represented by multiple clusters (average $N_{\text{cluster}}$ = 32–84, for early rounds; see Figure S1) with reproducibility improving with the use of random barcode sequences and a bottlenecking step ($N_{\text{cluster}}$ = 57–191, for rounds see Figure S1).

Data Fitting

Binding curves for MS2 protein binding (at protein concentrations 1–3000 nM) were collected for all clusters (e.g. **Fig. 1e**) in the presence and absence of the relevant small molecule ligand (e.g., 0 and 200 µM FMN). Fluorescence images were aligned to the sequencing data and individual cluster intensities were quantified with a custom analysis pipeline, normalized, and fit to a binding curve. Experimental reproducibility was determined by including a control set of one thousand standard designs in each Round, which exhibited excellent agreement between replicates (Fig S1).

The magnitude of switching of the designs was determined by the fold-change in the effective dissociation constant, $K_D$, for MS2 protein reporter binding in response to small-molecule ligand binding. The binding should be strong (low $K_D$) in the ON-state, where the RNA should display the reporter-binding motif, and weak (high $K_D$) in the OFF-state without the motif formed. In contrast to an activation ratio measured at a fixed reporter concentration, the fold change in $K_D$ is derived from all available data, measures changes from the full range of ON-state reporter affinities, and can be directly related to biophysical models. Furthermore, the fold change in $K_D$ gives the change in activation at lowest input ligand concentrations, where it should be maximal[10].

Data analysis

For each image, the cluster positions were aligned to the sequencing data coordinates and quantified using a custom analysis pipeline, as previously described[5]. The green (MS2) signal was normalized to the red (RNA) signal for each RNA cluster. The fluorescence signals for MS2-only controls were fit to a binding curve

$$F([MS2]) = F_{max}\frac{[MS2]}{[MS2]+K_D} \qquad (1)$$

Where $F_{\max}$ is the maximum fluorescence intensity, [MS2] is the MS2 concentration and $K_D$ is the observed dissociation constant. All control clusters were fit individually and the median $F_{\max}$ was then used to further normalize all cluster signals. These signals were then fit to the same binding curve as above and bad clusters were filtered out and the median $K_D$ and $F_{\max}$ values for each design were used for further analysis and for reporting data back to Eterna participants.

Scoring of participant designs

Full experimental data reports were returned to the Eterna community for each tested design. In addition, to aid in tracking progress, each experimentally tested Eterna design was given a single summary "Eterna Score" (on a scale of 0–100), based on sum of three subscores, as follows.

The Switch Subscore (*SS*) was calculated as

$$SS = 40\ Max\left(0, Min\left(1, log\ (FC)\ /log\ (FC_{high}\right)\right)$$

Here *FC* is the activation ratio (originally termed 'fold-change'), the ratio in the observed MS2 dissociation constant, $K_{D,\ obs}$ , for the design and $FC_{high}$ is an upper bound that represents the maximum signal. For the MS2 switches in this study, we originally set $FC_{high} = 26$. This value corresponded to the maximum activation ratio achievable based on estimates of the dissociation constant of the FMN aptamer (which were overestimated), FMN ligand concentration, and a desired $K_{D,\ obs}$ in the ON state of within 2-fold of the dissociation constant for the MS2 protein to a perfect aptamer ($K_{D,obs,ON} < 2\ K_{D,MS2}$). (To avoid confusion in scoring, we maintained the value of $FC_{high} = 26$ even as we improved measurements of the FMN aptamer dissociation constant; instead we noted to players in puzzle descriptions that activation ratios as high as 100 should be achievable and sought.)

The Baseline Subscore (*BS*) was calculated as

$$BS = 30\ Max\left(0, Min\left(1, 1.5 - 0.25\frac{K_{D,obs,ON}}{K_{D,MS2}}\right)\right)$$

where $K_{D,obs,ON}$ is the observed dissociation constant in the ON-state and $K_{D,MS2}$ is the affinity for the native MS2 hairpin, i.e., the score is maximized for $K_{D,obs,ON} < 2\ K_{D,MS2}$ and then decreased linearly with $K_{D,obs,ON}$ until being 0 for $K_{D,obs,ON} > 6\ K_{D,\ MS2}$. This subscore penalized switches whose ON configurations bound MS2 coat protein significantly worse than and isolated MS2 RNA hairpin. The value of $K_{D,\ MS2}$ was measured through control MS2 hairpin RNA sequences included in every experiment and fell in the range of 1-2 nM.

The Folding Subscore ($FS$) was

$$FS = 30\ Max\left(0, Min\left(1, \frac{(F_{max} - 0.3)}{0.4}\right)\right)$$

such that it was 0 for $F_{\max} < 0.3$, 30 for $F_{\max} \geq 0.7$, and linearly increasing in between. This subscore penalized RNA sequences that misfolded in their ON states and prevented MS2 protein binding.

Optimal activation ratio set by equilibrium thermodynamics

Throughout our study, we compared experimentally achieved activation ratios (AR; originally termed fold-changes, FC) with thermodynamically optimal ARs. The optimal AR of a riboswitch depends upon the affinity of the aptamer for the input ligand and the concentration of input ligand used to trigger the switch. A detailed description of this bound is given in reference[7], but for completeness, a brief derivation is given here.

As an example, we consider an FMN-controlled OFF-switch riboswitch construct controlling the formation of an MS2-binding hairpin aptamer, the expression platform used for the array riboswitches in this work. The thermodynamic efficiency of this construct can be estimated as follows. The partition function of the system is given by partitioning the system's full conformational ensemble into four states with the following conformations and energies ($E$):

| State | FMN aptamer formed | FMN bound | MS2 hairpin formed | MS2 bound | E |
|---|---|---|---|---|---|
| **1** | y | y | n | n | $-k_B T \ln\left(\frac{[FMN]}{K_{D,FMN}}\right)$ |
| **2** | y | n | n | n | 0 |
| **3** | n | n | y | n | $\Delta G$ |
| **4** | n | n | y | y | $\Delta G - k_B T\ ln\left(\frac{[MS2]}{K_{D,MS2}}\right)$ |

Here, $k_B$ is Boltzmann's constant, $T$ is the absolute temperature, [$FMN$] and [$MS2$] are the experimental concentrations of FMN and MS2, respectively, and $\Delta G$ is an unknown energy difference between the states without bound FMN or MS2. The fluorescent signal is only observed in state **4** and the probability of being in this state, derived from the partition function, is

$$p(signal) = \frac{[MS2]}{([MS2]+K_{D,obs})},$$

where

$$K_{D,obs} = K_{D,MS2}\left(1 + e^{\frac{\Delta G}{k_B T}}\left(1 + \frac{[FMN]}{K_{D,FMN}}\right)\right)$$

Accordingly, the normalized signal looks identical to a canonical binding curve. For ON-switches, a similar five-state model yields an analogous functional form, with

$$K_{D,obs} = K_{D,MS2} \frac{\left(1 + e^{\frac{\Delta G}{k_B T}}\right)\left(1 + \frac{[FMN]}{K_{D,FMN}}\right)}{\left(1 + e^{\frac{\Delta G}{k_B T}} + \frac{[FMN]}{K_{D,FMN}}\right)}.$$

For the constructs evaluated in this work, the reported ARs represent the ratio between the observed dissociation constants, $K_{D,obs}$, at two experimental ligand concentrations (for instance, 0 and 200 µM for FMN). In either ON or OFF switches, this ratio limits to $1 + \frac{[FMN]}{K_{D,FMN}}$ as the Boltzmann-weighted term involving $\Delta G$ becomes large. We may see this in the following expression for the AR of an OFF-switch:

$$\frac{K_{D,obs}([FMN]=200)}{K_{D,obs}([FMN]=0)} = \frac{1+e^{\frac{\Delta G}{k_B T}}\left(1+\frac{[FMN]}{K_{D,FMN}}\right)}{1+e^{\frac{\Delta G}{k_B T}}} < 1 + \frac{[FMN]}{K_{D,FMN}} = AR_{max}.$$

The expression for an ON-switch follows similarly. We refer the reader to ref.[7] for a more complete analysis of this thermodynamic model.

Independent assessment of an MS2 riboswitch using flow cytometry

As an independent test of an FMN-MS2 riboswitch discovered through RNA-MaP measurements, we displayed the RNA on particles to enable characterization of bound MS2 through flow cytometry. The sequence used was:

**BC_ JL-sng2-3.09 (underline is the active region)**

GGGUAUGUCGCAGAAGUAGCUAUCGGAGGAUAUUCAUACCGGAAACGGACAUGAGGAUCACCCA
UGUGGCGAAAGCCUUGGGAGAAGGCUGAUAGCUACAACUGCGACAUACCCAAAAAAAAAAAAAA
AAAAAAAAAAA

**Primers used for BC_ JL-sng2-3.09 assembly**

| | |
|---|---|
| BC_ JL-sng2-3.09_P1 | CGACAGCAGTTCTAATACGACTCACTATAGGGTATGTCGCAGAAGTAGCTATC |
| BC_ JL-sng2-3.09_P2 | GTGATCCTCATGTCCGTTTCCGGTATGAATATCCTCCGATAGCTACTTCTGCGACATACC |
| BC_ JL-sng2-3.09_P3 | GGAAACGGACATGAGGATCACCCATGTGGCGAAAGCCTTGGGAGAAGGCTGATAGCTACA |
| BC_ JL-sng2-3.09_P4 | TTTTTTTTTTTTTTTTTTTTTTTTTGGGTATGTCGCAGTTGTAGCTATCAGCCTTCTCCC |

**BC_MG-D3 (control sequence containing same flanking sequence)**

GGGUAUGUCGCAGAAGGGACACAAUGGACGCAGAUAAUCGGAUUCCCGACUGGCGAGAGCCAGG
UAACGAAUGGGCCGGUUUCUGUGUUAACGGCCGAACUGCGACAUACCCAAAAAAAAAAAAAAAA
AAAAAAAAA

**Primers used for BC_MG-D3 assembly**

| | |
|---|---|
| BC_JL_P1 | CGACAGCAGTTCTAATACGACTCACTATAGGGTATGTCGCAGAAGGGACACAATGGACGC |
| BC_JL_P2 | CATTCGTTACCTGGCTCTCGCCAGTCGGGAATCCGATTATCTGCGTCCATTGTGTCCCTT |

| | |
|---|---|
| BC_JL_P3 | GAGAGCCAGGTAACGAATGGGCCGGTTTCTGTGTTAACGGCCGAACTGCGACATACCCAA |
| BC_JL_P4 | TTTTTTTTTTTTTTTTTTTTTTTTTGGGTATGTCGCAGTTCGGCCGTTAACA |

DNA primer oligos were ordered from Integrated DNA Technologies (IDT) and full-length DNA oligos were assembled using standard PCR assembly protocols available at primerize.stanford.edu[11]. Briefly, 100 µL of 1X PCR mix containing Phusion DNA polymerase (Thermo Fisher Scientific) was prepared with 2 µM of primers P1 and P4, and 40 nM of primers P2 and P3. Then, 20 cycles of PCR were run at 98 °C (15 s) – 57 °C (30 s) – 72 °C (30 s). The correct length products were verified on agarose gels and transcribed using a Transcriptaid T7 High Yield transcription kit (Thermo Fisher Scientific) using the manufacturer's protocol. After DNase treatment for 30 minutes at 37˚C, RNA was purified using the RNA Clean & Concentrator kit (Zymo Research), resuspended in DEPC-treated $H_2O$ and the RNA concentration was determined using a UV spectrophotometer (NanoDrop, Thermo Fisher Scientific).

We prepared RNA on beads for flow cytometry as follows. The following oligo was ordered from the Protein and Nucleic Acid (PAN) Facility at Stanford University: /5`C6AmM/TTTTT TTTTT TTTTT TTTTT TTTT /3`InvdT/. This poly-T oligo contains an amino modification with a 6-carbon spacer on its 5' end, and was terminated using a 3' inverted deoxythymidine. 100 µL of 2.8 µm carboxylic acid beads (Dynabeads M-270, Invitrogen) were washed once with 100 µL 0.01 M NaOH and twice with 100 µL $H_2O$. The beads were incubated overnight with rotation at room temperature in 1X conjugation buffer (200 mM NaCl, 1 mM imidazole), 0.01% sodium dodecyl sulfate, 250 mM 1-ethyl-3-(3-dimethylaminopropyl) carbodiimide (EDC) and 100 µM of amino-modified poly-T oligo in 50% v/v DMSO. After incubation, the beads were washed twice with TT buffer (50 mM Tris-HCl pH 7.5, 0.5% Tween-20) and stored in 100 µL TNaTE buffer (10 mM Tris-HCl pH 7.5, 140 mM NaCl, 0.1% Tween-20, 1 mM EDTA) at 4 °C. 7 µL of poly-T-coated beads from above were added to 20 µL PBSMT (1X PBS, pH 7.2 (Gibco), 5 mM $MgCl_2$, 0.01% Tween-20) containing 2 µM of BC_JL RNA or BC_MG-D3 (control) and incubated at 37˚C for 5 minutes and then put on ice for 5 minutes. The buffer was removed and the beads were resuspended in 360 µL PBSMT. 500 uL of 2.4x solutions of FMN (1.944 mM) and SNAP-549-labelled MS2 coat protein (MCP, 1.944 µM) were prepared in PBSMT and these underwent 1:4 dilutions a total of five times. Assays were prepared using 25 µL of each 2.4x solution or PBSMT, as well as 10 µL of RNA-coated beads (to give a total volume per tube of 60 µL) such that the final concentrations ranged from 810 µM to 3.2 µM and 0 µM for FMN and from 810 nM to 3.2 nM and 0 nM for MCP. Each sample was analyzed using a Sony SH800S Cell Sorter and data for 10,000 events were collected per sample. Beads were excited using a 561 nm laser and their emitted fluorescence was measured from the 600±60 nm emission channel.

**Fluorescence measurements for light-up sensor challenges**

Small molecules for light-up sensor challenges

We developed Eterna puzzles that incorporated previously-described aptamers against the small molecules theophylline (Theo), flavin mononucleotide (FMN), tryptophan (Trp), and tetracycline (Tet), as inputs; and malachite green (MG) and 3,5-difluoro-4-hydroxybenzylidene imidazolinone (DFHBI), as outputs. All molecules were purchased from Sigma-Aldrich unless otherwise specified. The malachite green aptamer (MGA) and DFHBI aptamer (RNA Spinach) belong to a class of light-up RNAs that effect >1000-fold fluorescence enhancement upon binding the target fluorophore molecule. Specifically, our puzzles coupled small molecule (input) binding to turn-on fluorescence activation for the following input/output pairs: Theo/MG, FMN/MG, Trp/MG, Trp/DFHBI, Tet/MG, and Tet/DFHBI. Pilot experiments with FMN/DFHBI puzzles showed significant fluorescent overlap between FMN and DFHBI that precluded characterization, and these were not continued. In addition, experiments with Theo/DFHBI and ATP/DFHBI sensors showed interference of the aromatic input ligands with RNA Spinach, likely through competitive binding with DFHBI, and these were also not pursued.

In each light-up sensor round, we collected 5-6 participant designs from two puzzles at a time. The characterized designs were chosen by participants in a weekly community-wide vote, appended with complementary flanking sequences, ordered as DNA oligos (Integrated DNA Technologies) designed with Primerize[11], PCR amplified using standard protocols, transcribed using a TranscriptAid T7 High Yield Transcription Kit (ThermoFisher), and purified using an RNA Clean & Concentrator-25 kit (Zymo Research) using the manufacturer protocols.

Light-up fluorescence assays

Assays were prepared in 96-well, half-area, flat-bottomed plates (Costar). Each participant design was titrated at 8 different concentrations of their respective light-up dye both in the absence and presence of their target small molecule (see table below for input ligand concentrations). The light-up dyes were titrated from 10 $\mu$M to 16 nM in an 8-point, 1:2.5 dilution series, except for in the first light-up series where the dye was titrated from 5 $\mu$M to 40 nM in an 8-point, 1:2 dilution series. The final RNA concentration in each well was 200 nM at a final volume of 100 $\mu$L. Concentrations for small molecules were obtained from their extinction coefficients:

| Molecule | $\lambda_{max}$ | $\epsilon\ (cm^{-1}M^{-1})$ | Concentration |
|---:|---|---:|---|
| **Theophylline** | 277 | 10,200 | 1.2 mM |
| **Tryptophan** | 278 | 5,579 | 2.4 mM |
| **Tetracycline** | 366 | 14,150 | 60 $\mu$M |
| **FMN** | 450 | 9,910 | 200 $\mu$M |
| **DFHBI** | 405 | 11,864 | varied |
| **MG** | 617 | 148,900 | varied |

After equilibration of the wells for 30 minutes at room temperature, fluorescence intensity for each well was measured using a Tecan Infinite 200 PRO plate reader. For switches containing the Spinach aptamer to DFHBI, excitation was at 466 $\pm$ 9 nm and emission measured at 508 $\pm$ 20 nm. For switches containing the MGA aptamer to malachite green, excitation was at 625 $\pm$ 9 nm and emission measured at 660 $\pm$ 20 nm.

Data Fitting

The resulting fluorescence intensities in each well were plotted against the light-up (LU) dye concentration and fit to the equation

$$B = B_{max} * \frac{[LU]}{[LU] + K_D^{LU}}$$

Where B is the brightness at each concentration [LU] of the respective light-up dye, $B_{max}$ is the brightness at saturation for the given sample, and $K_D^{LU}$ is the dissociation constant for the design to the dye. To determine experimental activation ratios, we determined the ratio of LU fluorescence without and with ligand at lowest LU ligands, analogous to the RNA-MaP measurements above. More specifically, for each design, we determined the $B_{max}$ and $K_D$ in both the absence (-) and presence (+) of the input ligand. We determined the activation ratio of each design by comparing the signal in the presence and absence of input ligand:

$$AR = \frac{B^{(+)}}{B^{(-)}} = \frac{B_{max}^{(+)}}{B_{max}^{(-)}} * \frac{[LU] + K_D^{LU(-)}}{[LU] + K_D^{LU(+)}}$$

Which, in the case for an ON switch where $K_D^{LU(-)} > K_D^{LU(+)}$ approaches its maximum at low [LU]:

$$AR^{LU \to 0} = \frac{B_{max}^{+SM}}{B_{max}^{-SM}} * \frac{K_D^{LU(-)}}{K_D^{LU(+)}}$$

The activation ratio of each design was then compared to its theoretical max

$$AR_{max} = \frac{[input]}{K_d} + 1$$

Where the $K_d$ value of the switch to the respective small molecule was determined using DMS Chemical Mapping under identical buffer conditions and using a non-switching ("always ON") ligand-binding control sequence of similar architecture to the respective puzzle.

Data Analysis

The $K_D^{LU}$ and $B_{max}$ values were computed for each switch design in the presence or absence of the respective small molecule. Unlike the MS2 switches characterized on the array, the $B_{max}$ value for a given RNA can change based on the predominant state of the switch. That is, two states of the same switch can promote different levels of fluorescence enhancement of the LU dye independently of their affinity to the dye. For this reason, we report three ratios ($K_D^{LU}$, $B_{max}$, and $AR^{LU \to 0}$) for each participant design based on their fit in the absence or presence of the relevant small molecule ligand. Switches whose $K_D^{LU}$ were higher than the limits of detection or whose $B_{max}$ values did not significantly exceed background (i.e., designs where both fitted $B_{max}$ values were >20-fold lower than the $K_D^{LU}$ of the non-switching control light-up aptamer used in each round) were marked as non-functional.

Kinetic characterization of light-up sensors

We wished to determine the rate-limiting step for switching of sequence BH_E029, a Trp/MG "ON" switch through kinetic measurements. We characterized the sequence

```
BH_E029:
GGGAGAGUGGCAGCUUACAACGUAACGAUGCGGCCGCCACUGAUCCGACUGGUUAACAGAAAGUCCUGACCAGGUAA
CGAAUGGUCAGGACCGGGCAUUGUUACGAAGUGGGCUGUUGCUUUUCC
```

Stock 3x solutions of MG (172 nM), RNA (150 nM), and Tryptophan (7.2 mM) were prepared in 1X PBS with 5 mM $MgCl_2$ at pH 6.1 (PBSM). To prepare the pre-equilibrated controls, 30 minutes prior to injection we mixed 30 µL each of 3x MG, 3x RNA, and either 3x Trp (yellow curves) or 1x PBSM (red curves) buffer. To prepare the non-equilibrated controls, we prepared the same mixtures except withholding either RNA (**Fig. S4A**), or Trp (**Fig. S4B**). At the time of measurement, all wells had final reagent concentrations of 64 nM MG, 50 nM RNA, and 2.4 mM Trp in a total volume of 90 µL and all wells were prepared in duplicate.
To determine the rate at which Trp bound to the switch (Figure A), we injected 30 µL of 3x RNA into both of the Trp(+) and Trp(−) wells to yield the final concentration given above and measured the fluorescence intensity of these 4 samples and their pre-equilibrated controls every ~3.3 s for 30 minutes. We then fit the fluorescence ($B$) as a function of its maximum fluorescence ($B_{max}$) time ($t$) and the association ($k_{on}$) :

$$B = B_{max} * \left(1 - exp\left(-k_{on} \cdot (t_0 + t)\right)\right)$$

Where B represents the fluorescence intensity at each point, $B_{max}$ the maximum brightness at equilibrium, $k_{on}$ the association rate ($s^{-1}$), $t$ the time from injection (seconds), and $t_0$ the delay (seconds) between injection and the first measurement (about 6 seconds).

To determine the rate at which MG bound to the switch (Figure B), we injected 30 µL of 3x MG into both of the Trp(+) and Trp(−) wells to yield the same final concentrations as above and measured the fluorescence intensity of these 4 samples and their pre-equilibrated controls every ~3.3 s for 5 minutes. The association rate of MG with the switch was too fast for measurement under these conditions.

**Chemical mapping measurements**
To test reversible structural toggling of RNA switches by FMN without MS2 coat protein, RNA molecules for the JL-sng2-3.09 sensor were generated by T7 RNA polymerase *in vitro* transcription and purified, as in our previous protocol [12]. This RNA was initially hybridized to magnetic poly-dT beads (Poly(A) Purist, Ambion) and incubated at 4 °C. Buffer and salt conditions were identical to those in the RNA-MaP experiments (100 mM Tris-HCl, pH 7.5, 80 mM KCl, 4 mM $MgCl_2$) and the FMN concentration was varied by repeatedly pulling down (via magnetic stand) and re-suspending the beads in buffer without or with FMN (0 or 200 µM). Incubation times after each buffer exchange was 5 min, and the experiments were carried out at ambient temperature (24 °C). Every three cycles, two sub-samples (20 µL each) from the large experimental volume (initially 1.28 mL) were collected and chemically probed using 1-methyl-7-nitroisatoic anhydride (1M7) or left unmodified, as controls. Workup of these sub-samples, including purification, reverse transcription, and quantitative analysis in HiTRACE [13], followed our previous protocol [12].

Measurements of affinities of each RNA aptamer to its target ligand (FMN, tetracycline, tryptophan, theophylline) was also carried out by 1M7 or DMS chemical mapping (see Table S3). The sequences used for these chemical mapping experiments included RNA constructs designed to present the aptamers within stable hairpins, flanking sequences that matched those of

RNA-MaP or light-up fluorescence experiments, as appropriate, and additional 3’ sequences used to prime reverse transcription, as follows:

| Aptamer | Condition | RNA Sequence |
|---|---|---|
| FMN | RNA-MaP | GGAAAGGCGUGU**AGGAUAU**GCUUCGGC**AGAAGG**ACACGCCAAAGAAACAACAACAACAAC |
| Tetracycline | RNA-MaP | GGAAAGGCG**CUAAAACAUACC**AGUUCGCU**GGAGAGGUGAAGAAUACGACCACCUAG**CGCCAAAGAAACAACAACAACAAC |
| Tryptophan | RNA-MaP | GGAAAGGCGAG**CGGCCGCCACU**GUUCGC**AGGACCGG**GCUCGCCAAAGAAACAACAACAACAAC |
| Theophylline | RNA-MaP | GGAAAGGCGAG**GAUACCAG**CGUUCGCG**CCCUUGGCAGC**CUCGCCAAAGAAACAACAACAACAAC |
| FMN | Light-up | GGGAACGUCUCCGAGUAGGAGACGAAGGGAGAGUGGCAGCUUACAAUUCCAGCCGACCACUGAAAUACUCUAG**AGGAUA**UGACAGACAAAGUCUGUC**AGAAGG**CUAGAGCAGUGGGUAACGAAUGCUGGAAAAGUGGGCUGUUGCUUUUCCAAGCUUAUCGAGUAGAUAAGCAAAAAGAAACAACAACAACAAC |
| Tetracycline | Light-up | GGGAACGUCUCCGAGUAGGAGACGAAGGGAGAGUGGCAGCUUACAACG**CUAAAACAUACC**UGCACCACCGACCUUGAAAACAAGGGUAACGAAUGUGGAGCA**GGAGAGGUGAAGAAUACGACCACCUAG**CGAAGUGGGCUGUUGCUUUUCCAAGCUUAUCGAGUAGAUAAGCAAAAAGAAACAACAACAACAAC |
| Tryptophan | Light-up | GGGAACGUCUCCGAGUAGGAGACGAAGGGAGAGUGGCAGCUUACAAACUACAGUUG**CGGCCGCCACU**GCGAAGGACGGGUCCCUCGAAAGAGGUUGAGUAGAGUGUGAGCGC**AGGACCGGG**CAACAGUAGUAAGUGGGCUGUUGCUUUUCCAAGCUUAUCGAGUAGAUAAGCAAAAAGAAACAACAACAACAAC |
| Theophylline | Light-up | GGGAACGUCUCCGAGUAGGAGACGAAGGGAGAGUGGCAGCUUACAACCUGAAGGACGGGUCCCCAGUAAUAAU**GAUACCAG**GAACAAAAGUUC**CCCUUGGCAGC**AUCUGGGUUGAGUAGAGUGUGAGCAGGAAGUGGGCUGUUGCUUUUCCAAGCUUAUCGAGUAGAUAAGCAAAAAGAAACAACAACAACAAC |

### Supplementary Text

Features of top-performing designs

To understand how top participants achieved success in designing efficient standalone riboswitches, the community engaged in ‘all-hands’ online discussions, presentations during annual Eterna conferences (EternaCon), and compilation of puzzle-solving strategies in game forum posts and widely shared Google documents. As expected, participants approached new puzzles in a highly varied manner, but several recurring principles were articulated:

First, no single puzzle architecture was universally suitable for all given input-output combinations. Rather, it was necessary to test various puzzle architectures that differed in the relative placement of the input and output aptamers. For the first FMN-MS2 ON-switch puzzles (main text Figure 2), many successful solutions placed the FMN input aptamer and the MS2 output hairpin as close as possible to each other, and sequestered other nucleotides available for design into ‘static stems’. Indeed, several independent solutions converged on an architecture that was identical in its ordering of the FMN aptamer segments, MS2 segments, and static stems up to circular permutation of the sequence (compare three panels of main text Figure 2G). At the same time, the light-up sensor challenges involved larger aptamer sequences than the FMN aptamer, and both the aptamer for the input ligand and the aptamer for the output fluorophore involved two segments that needed to be placed. These constraints necessitated use of larger segments of the available sequence for switching and obviated the sequestering of extra nucleotides into static stems. Nevertheless, new rules were proposed for these more difficult challenges. While the most successful participant designs all included a ‘split’ aptamer (where one aptamer is nested inside the other), determining which aptamer was split to yield optimal performance was often determined by considering the affinity of the aptamer (the higher affinity aptamer was generally split), the individual aptamer sequences (aptamers predicted to fold stably were often split), and any sequence similarity that may exist between the two aptamers.

Exceptions to these general rules are also apparent, highlighting that optimal switch design requires testing not only multiple switch designs, but also multiple switch architectures.

Second, in initial FMN-MS2 challenges that allowed for iterative refinement of solutions, participants commonly modified the most successful designs in every round in order to improve performance in subsequent rounds. This is evident as the emergence of design "families" sharing high sequence similarity (see Fig 2B-C). In addition, the most successful designs were confirmed to switch *in silico* using multiple folding engines. While the Vienna RNA 1.8.5 folding package was originally the de facto RNA folding engine for Eterna, veteran Eterna participants suggested that other engines such as Vienna 2.0 and NuPACK showed higher predictive power, and that the best switches were predicted to function in all three engines. For this reason, we provided participants in-game tools to model their switches in multiple folding engines in all later rounds of puzzles. At the same time, we note that none of these modeling packages provide strong predictive power in modeling actual activation ratios (see, e.g., ref. [9]), underscoring the need for design principles and architectures that are robust to errors in these packages.

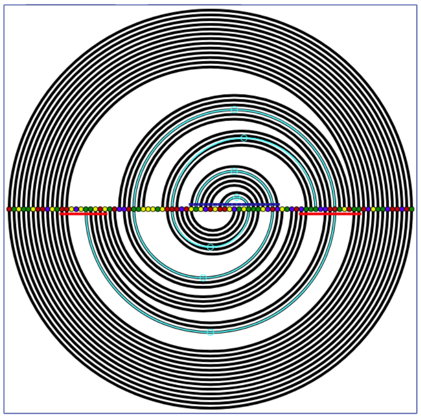

Example arc plot depicting a spiral (in light blue), indicating that bases have a high likelihood of switching base-pairing between the ligand-free and ligand-bound state. From the document "Arc-Plot Tool Intro" by Eterna players jandersonlee and Eli.

Third, some participants showed adaptability and ingenuity when approaching these challenges. As one example, several participants independently began incorporating multiple input aptamers into their switch designs; thus 'cheating' the expected thermodynamic maximum activation ratio for a single-input switch (Supplemental Figure S3 and ref. [10]).

Last, several participants spearheaded the development and dissemination of tools that granted new insights into individual switch puzzles and/or solutions. During light-up challenges, players developed visualization tools for simultaneously tracking both states of the sensor and tabulating metrics that were not provided in the Eterna interface. For example, one metric was a solution's 'x-ratio'- or ratio of predicted pairing probabilities, in the absence versus presence of input ligand, of the outermost base-pairs of the split aptamer, which approximately predicts the actual activation ratio of the design. Participants generally sought a high predicted x-ratio, and this feature was incorporated into a participant-designed 'arcplot' online tool and distributed amongst the community. Arcplots show pairing probabilities of all bases in the absence and presence of ligand in a single chart. Participants also disseminated screenshots of arcplots with striking features (e.g., spirals).

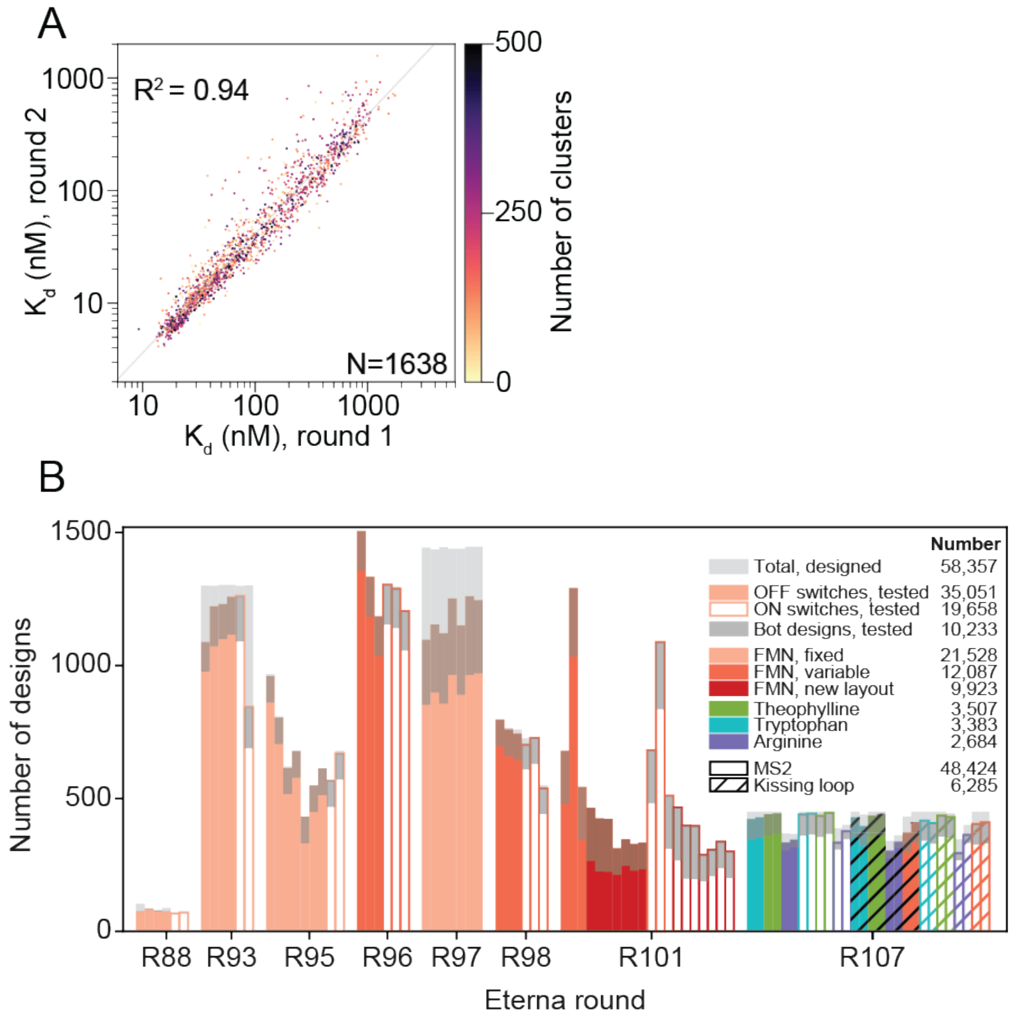

**Fig. S1. Reproducibility and scale of RNA-MaP experiments.** A) Reproducibility of a set of standard sequences spanning the range of detection. The replicates differ by time, protein preparation, DNA synthesis, and experimentalist. B) Number of Eterna designs created and tested. In total, 58,357 RNA switches were designed by the Eterna community, and the majority of these were successfully tested on the RNA array over seven Eterna rounds. The switches could either unfold the MS2 hairpin and turn off MS2 protein binding upon addition of ligand (OFF switches) or turn it on (ON switches). In each round, a minority of solutions were generated by computer algorithms (Ribologic). Three puzzle layouts were used for FMN. In the first (FMN, fixed), the position of both the FMN aptamer and the MS2 hairpin were fixed. In the second (FMN, variable), the MS2 hairpin could be placed in any location not overlapping the required aptamer residues. In the third (FMN, new layout), the position and direction of the aptamer arms as well as the sequence length in between them were changed. In the last round (R107), puzzles with new aptamers (Theophylline, Tryptophan, and Arginine) were designed, using either the previous MS2 reporter or a new Kissing loop reporter system.

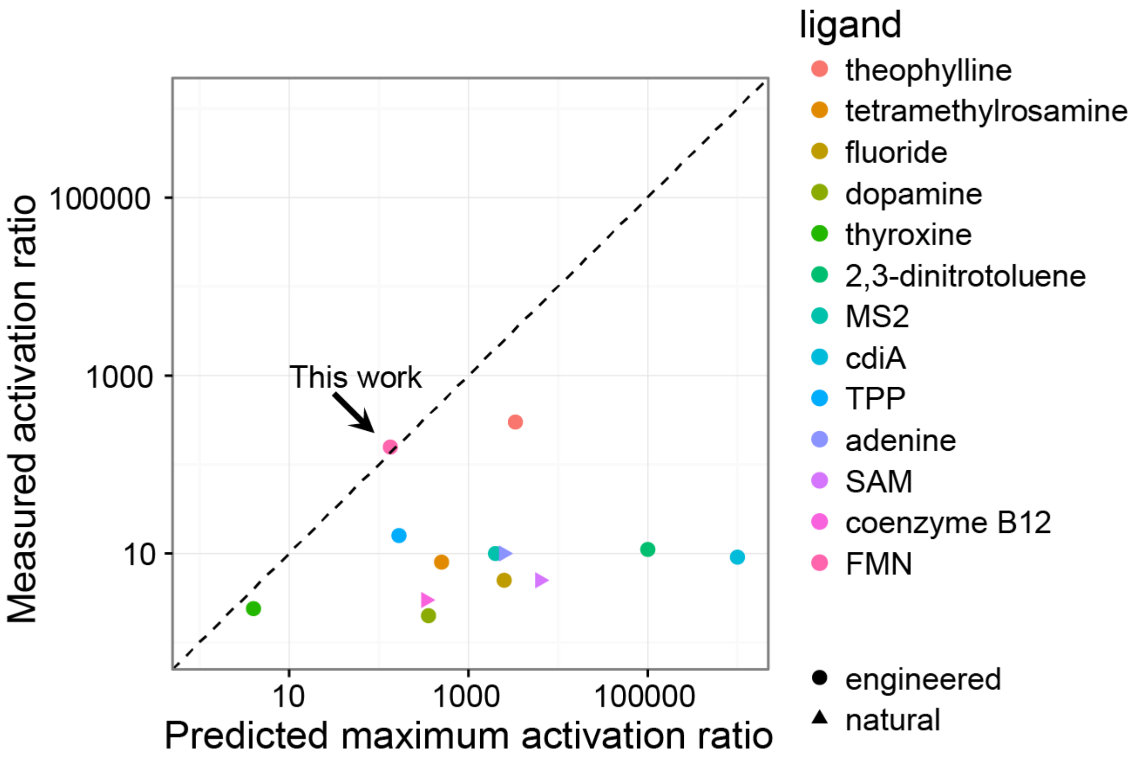

**Fig. S2. Predicted and measured activation ratios for riboswitches in the literature.** Typically, the activation ratio (*AR*) is ~10-fold or below, less than the maximum predicted from the affinity of the ligand and the concentrations used in the experiments. The FMN/MS2 riboswitches in this work (arrow) reach activation ratios nearly equal that of the maximally predicted value, calculated from the measured effective binding constants ($K_D$) and measured affinities for FMN (by chemical mapping) and MS2 (measured for controls on the RNA array in the same experiment).

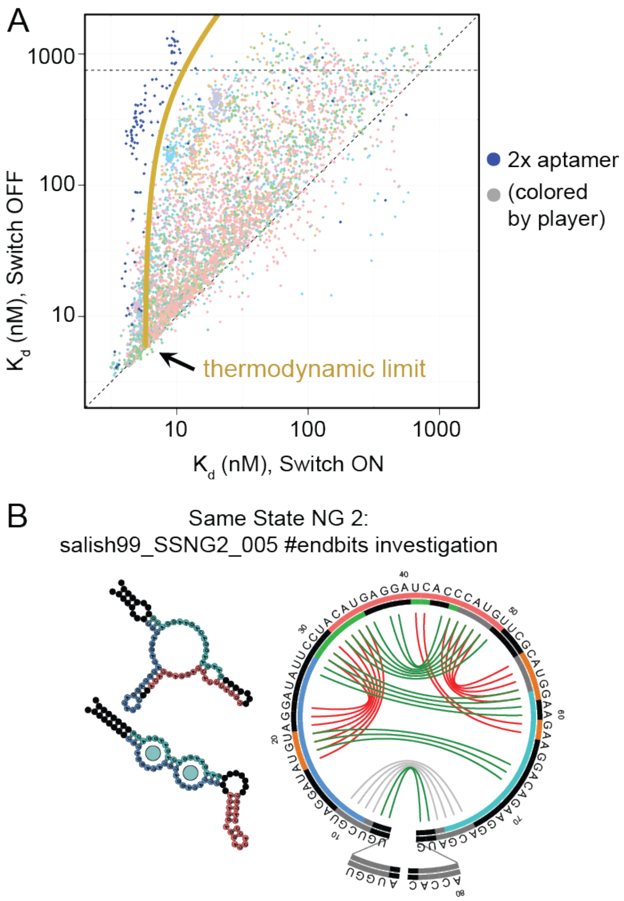

**Fig. S3**. **Cheat solutions with double input aptamers can surpass the thermodynamic limit for single input aptamers.** A) $K_d$ values for both states of "Same State NG 2" puzzle designs. The predicted thermodynamic limit for the $K_d$ in the OFF-state, based on the $K_d$ in the ON-state, is shown in green. Player solutions (colored by player as in main text Fig. 2E) are shown, with solutions that use two FMN aptamer sequences colored in blue. These designs often exceed the calculated thermodynamic limit (gold curve). These "cheat" solutions exhibit high OFF-state $K_d$ values while maintaining small $K_d$s in the ON-state. B) Predicted secondary structures for a well-performing double aptamer solution with predicted invariant base pairs (grey segments) and base pairs in the absence (red) or presence (green) of the FMN ligand. Outer circles display the MS2 hairpin and complementary segments in color; inner circle displays the FMN aptamer and complementary segments (as in main text Fig 2F).

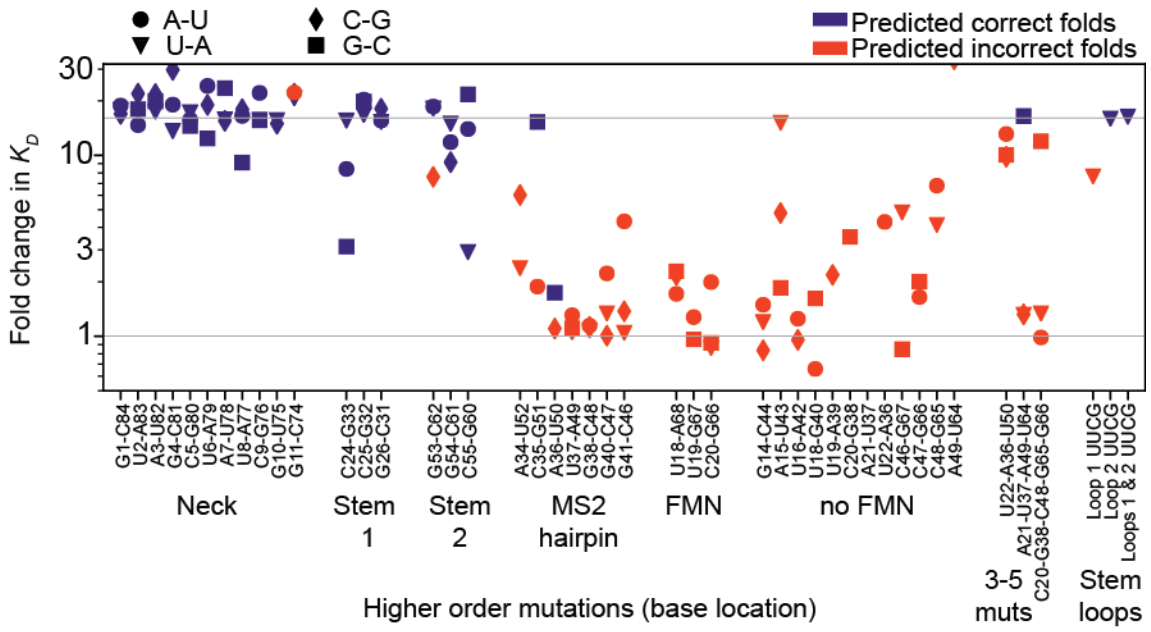

**Fig. S4. Testing structural mechanisms by mutagenesis and functional measurements.** Fold-change in $K_d$ (activation ratio) for double and higher order mutants of the best Same State NG 2 puzzle solution (see table S2). Identical data as in Figure 2G but with all mutations indicated. All base pairs in at least one of the two secondary structures were mutated, changing the bases at that location to each of the other three canonical base pairs. Mutated sequences predicted to fold into the same secondary structures in the FMN and no FMN states are shown in blue. Most of these are located in the neck or the preserved stems. The other mutations, including those of the MS2 hairpin and the base pairs flanking the FMN aptamer, are predicted to break at least one of the two structures and are plotted in red. On the right are sequences with three or more base changes that preserve base pairing by propagating mutations or by mutating the stem tetraloops.

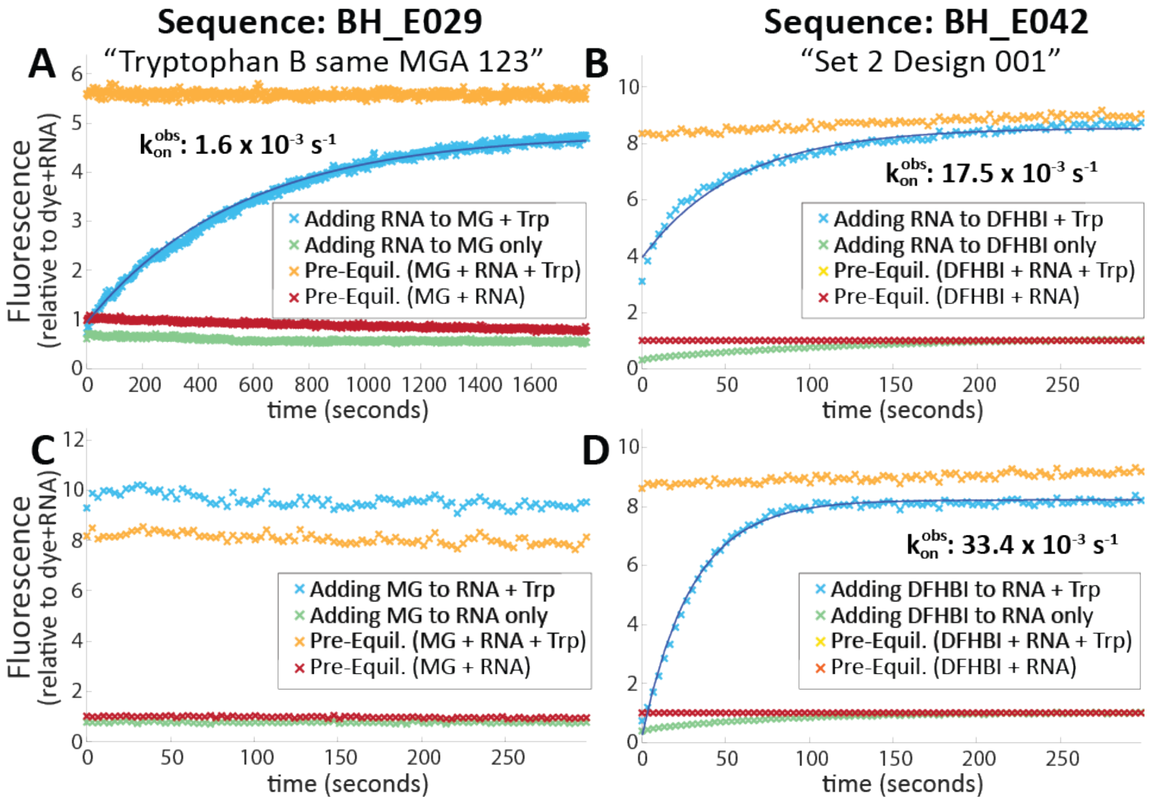

**Fig. S5. Testing kinetics of ligand-triggered switching.** Adding RNA into a solution of pre-equilibrated MG (A) or DFHBI (B) containing tryptophan (Trp) leads to variable rates of increasing signal (blue curves) approaching that of the same solution pre-mixed 30 minutes earlier (orange curves). The variable Trp association rates between these sensors—which contain identical Trp-binding aptamer motifs—suggests that structural rearrangement is the major energetic barrier for design BH_E029. In contrast, injecting MG (C) or DFHBI (D) into a solution of *pre-equilibrated* RNA + Trp (blue curve), we see that binding of the dye molecule happens rapidly, quickly approaching (or surpassing) the fluorescence signal of the same sample pre-mixed 30 minutes earlier (yellow curve). These data demonstrate switching kinetics on the order of 10s of seconds (for BH_E042) or minutes (BH_E029) with the slower switch likely gated by timescales of structural rearrangement. In all experiments, little fluorescence is detected for the analogous assay without any tryptophan (red and green curves), as expected for Trp-triggered light-up sensors.

| | | Switch Design Round (Eterna Round) | | | | | | |
|---|---|---|---|---|---|---|---|---|
| | **Puzzle name** | **1 R88** | **2 R93** | **3 R95** | **4 R96** | **5 R97** | **6 R98** | **7 R101** |
| | **OFF switches** | | | | | | | |
| 1. | Exclusion 1 | 75/104 | 1089/1301 | 960/968 | | 1097/1443 | | |
| 2. | Exclusion 2 | 85/87 | 1223/1300 | 804/809 | | 1154/1437 | | |
| 3. | Exclusion 3 | 77/81 | 1231/1303 | 615/620 | | 1121/1445 | | |
| 4. | Exclusion 4 | 75/88 | 1258/1303 | 678/680 | | 1252/1439 | | |
| 5. | Exclusion 5 | | | 432/432 | | 1151/1439 | | |
| 6. | Exclusion 6 | | | 549/549 | | 1261/1446 | | |
| 7. | Broourd's mod of Exclusion 4 | | | 613/613 | | 1245/1446 | | |
| 8. | Exclusion NG 1 | | | | 1505/1518 | | 796/797 | 679/679 |
| 9. | Exclusion NG 2 | | | | 1333/1336 | | 759/765 | 1291/1291 |
| 10. | Exclusion NG 3 | | | | 1185/1186 | | 743/759 | 543/544 |
| 11. | Inverted Exclusion NG 1 | | | | | | | 465/465 |
| 12. | Inverted Exclusion NG 2 | | | | | | | 425/426 |
| 13. | Inverted Exclusion NG 3 | | | | | | | 424/424 |
| 14. | Small Loop Exclusion NG 1 | | | | | | | 312/315 |
| 15. | Small Loop Exclusion NG 3 | | | | | | | 346/346 |
| 16. | Inverted Small Loop Exclusion NG 1 | | | | | | | 328/328 |
| 17. | Inverted Small Loop Exclusion NG 3 | | | | | | | 334/334 |
| | **ON switches** | | | | | | | |
| 18. | Same State 1 | 67/67 | 1262/1303 | 568/568 | | | | |
| 19. | Same State 2 | 70/75 | 845/1302 | 670/679 | | | | |
| 20. | Same State NG 1 | | | | 1304/1305 | | 702/726 | 682/682 |
| 21. | Same State NG 2 | | | | 1289/1296 | | 728/732 | 1088/1091 |
| 22. | Same State NG 3 | | | | 1205/1207 | | 538/550 | 511/511 |
| 23. | Inverted Same State NG 1 | | | | | | | 466/466 |
| 24. | Inverted Same State NG 2 | | | | | | | 398/398 |
| 25. | Inverted Same State NG 3 | | | | | | | 397/397 |
| 26. | Small Loop Same State NG 1 | | | | | | | 288/288 |
| 27. | Small Loop Same State NG 3 | | | | | | | 307/307 |
| 28. | Inverted Small Loop Same State NG 1 | | | | | | | 338/338 |
| 29. | Inverted Small Loop Same State NG 1 | | | | | | | 301/301 |

**Table S1.** Number of FMN/MS2 riboswitch sequences designed and successfully tested per puzzle and round.

| Puzzle Name (R107) | Max participant AR | Max Ribologic AR | Thermodynamic maximum AR |
|---|---|---|---|
| **Theo Exc A** | 27 | 3.9 | 92 |
| **Theo Exc B** | 29 | 9.92 | 92 |
| **Trp Exc A** | 40.32 | 4.29 | 198 |
| **Trp Exc B** | 15.31 | 3.2 | 198 |
| **Theo SS A** | 13.83 | 2.6 | 92 |
| **Theo SS B** | 34.97 | 15.37 | 92 |
| **Trp SS A** | 13.95 | 4.55 | 198 |
| **Trp SS B** | 24.60 | 2.96 | 198 |

**Table S2.** Top activation ratios for MS2-activating sensors from Eterna players and Ribologic automated design, compared to thermodynamic maximum expected based on ligand concentrations. Data are from challenges without multiple rounds of refinement. Abbreviations: Theo, theophylline; Trp, tryptophan; Exc, exclusion (OFF-switches, ligand binding disfavors MS2 binding); SS, same-state (ON-switches, ligand binding favors MS2 binding); AR, activation ratio.

| Aptamer | Conditions* | Mod | $K_d$ | $K_d$ error (+) | $K_d$ error (-) | Units | Assay Conc. |
|---|---|---|---|---|---|---|---|
| FMN | RNA-MaP | 1M7 | 1.71 | 0.60 | 0.44 | µM | 200 µM |
| Tetracycline | RNA-MaP | 1M7 | 622.07 | 132.12 | 105.63 | nM | 60 µM |
| Tryptophan | RNA-MaP | DMS | 12.09 | 10.50 | 5.19 | µM | 2.4 mM |
| Theophylline | RNA-MaP | DMS | 13.11 | 9.21 | 4.86 | µM | 1.2 mM |
| FMN | Light-up | DMS | 1.57 | 0.58 | 0.41 | µM | 200 µM |
| Tetracycline | Light-up | DMS | 57.50 | 22.58 | 15.88 | nM | 60 µM |
| Tryptophan | Light-up | DMS | 11.59 | 9.49 | 5.00 | µM | 2.4 mM |
| Theophylline | Light-up | DMS | 239.54 | 161.15 | 96.27 | µM | 1.2 mM |

* RNA-MaP solution conditions (100 mM Tris-HCl, pH 7.5, 80 mM KCl, 4 mM $MgCl_2$, 37 °C) and light-up sensor conditions (1x PBS, pH 6.1 (Gibson); 5 mM $MgCl_2$, 24 °C).

**Table S3.** Affinity constants as determined using dimethyl sulfate (DMS) or 1-methyl-7-nitroisatoic anhydride (1M7) chemical mapping.
